# Extracellular Matrix acts as pressure detector in biological tissues

**DOI:** 10.1101/488635

**Authors:** Monika E. Dolega, Benjamin Brunel, Magali Le Goff, Magdalena Greda, Claude Verdier, Jean-François Joanny, Pierre Recho, Giovanni Cappello

**Affiliations:** LIPhy, CNRS–UMR 5588, Université Grenoble Alpes, 38000 Grenoble, France; Collège de France, PSL Research University, 11 place Marcelin Berthelot, 75005 Paris, France; Ecole Superieure de Physique et de Chimie Industrielles de la Ville de Paris - ParisTech, 75005 Paris, France; Physico-Chimie Curie CNRS–UMR 168, Institut Curie, 5 Rue Pierre et Marie Curie, 75005 Paris, France

## Abstract

Imposed deformations play an important role in morphogenesis and tissue homeostasis, both in normal and pathological conditions ^1–5^. To perceive mechanical perturbations of different types and magnitudes, tissues need a range of appropriate detectors ^6–8^, with a compliance that has to match the perturbation amplitude. As a proxy of biological tissues, we use multicellular aggregates, a composite material made of cells, extracellular matrix and permeating fluid. We compare the effect of a selective compression of cells within the aggregate, leaving the extracellular matrix unstrained, to a global compression of the whole aggregate. We show that the global compression strongly reduces the aggregate volume ^9–13^, while the same amount of selective compression on cells has almost no effect ^14,15^. We support this finding with a theoretical model of an actively pre-stressed composite material, made of incompressible and impermeable cells and a poroelastic interstitial space. This description correctly predicts the emergent bulk modulus of the aggregate as well as the hydrodynamic diffusion coefficient of the percolating interstitial fluid under compression. We further show that, on a longer timescale, the extracellular matrix serves as a sensor that regulates cell proliferation and migration in a 3D environment through its permanent deformation and dehydration following the global compression.

## Introduction

There are several evidences that the mechanical environment of a cell plays a fundamental role on cell fate, both in terms of proliferation and differentiation. Discher et al.^16^ observed that stem cells differentiate into different cell types, when cultured on polyacrylamide gels of different stiffness covered with collagen. Trappmann et al.^17^ ascribe such a differentiation to the collagen/polyacrylamide cross-link density, which accidentally correlates with the gel stiffness. Nevertheless, both studies point to the crucial role played by the mechanical interaction between the cell and its microenvironment. This becomes particularly relevant in an tissue structure, where the cells are in contact with an ExtraCellular Matrix (ECM) which is essential to the structure and function of the tissue as cells respond to chemical and mechanical signals provided by the surrounding ECM, but also constantly remodel and/or produce it^18^. ECM biochemical composition and 3D organization varies from thin layers of basement membrane, in epithelial tissues ^19^, to more abundant layers, in between cells in tumors ^20,21^. Such abundant and crosslinked ECM leads to tissue stiffening, which in turn increases the risk for malignancy by enhanced integrin signaling ^20^. Yet, how the presence of the ECM in multicellular structures determines their mechanical behavior remains to be determined and could be a key factor in some cases. This is exemplified by the fact that the compressibility modulus of individual cells^15^ is several orders of magnitude higher than the one of multicellular aggregates of similar cells^22,23^. In this article, we use composite MultiCellular Spheroids (MCS) composed of murine colon cancer cells (CT26 WT) to design an experimental technique to either selectively compress the single cells in the spheroid or globally the whole MCS. Next, we propose a mechanical model that quantitatively predicts the emergent compressibility and permeability of the MCS in which impermeable and incompressible cells coexist with a soft poroelastic ECM (imaged by immunolabeling of fibronectin in Figure 1a) permeated by interstitial fluid. In parallel, we show that this peculiar rheology has a strong impact on cell proliferation and migration in MCS under compression.

**Figure 1:**
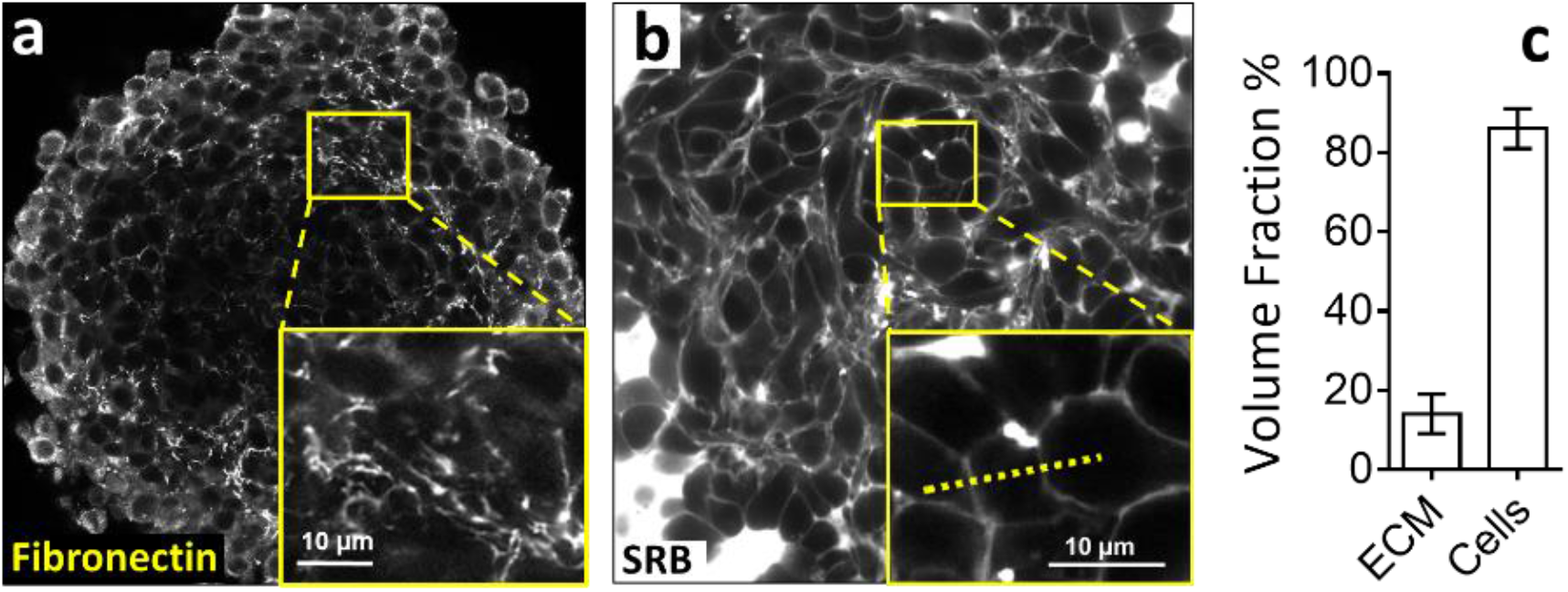
Extracellular Matrix volume fraction in multicellular spheroids. (**a**) Immuno-fluorescent staining of fibronectin and (**b**) intercellular space observed by confocal microscopy. The intercellular space is filled by sulphorhodamin-B, a hydrophilic colorant that does not penetrate the plasma membrane. (**c**) Taking into account the confocal resolution, we measure that the intercellular space has an average thickness 930±50nm. We estimate that the extracellular space represents 14%±5% of the total volume of a whole multicellular spheroid.

## Results

To quantify the ECM interstitial volume fraction *n_m_* in MCS, we performed negative imaging by supplementing the culture medium with sulphorhodamine B, a hydrophilic fluorophore that permeates the luminal space (Figure 1b). This approach allows for live confocal imaging and avoids the potential artefacts due to fixation and cryosectioning. Taking into account the confocal resolution (270 nm in our experimental conditions), we measured an average thickness of the intercellular space of 0.9±0.1 μm. Considering that CT26 cells have a typical diameter of about 20μm, we estimate an interstitial volume fraction of *n_m_* = 14±5% (Figure 1c and SI-1).

To understand the role of ECM in perceiving and transmitting mechanical perturbations we established a novel method that allows compressing either globally our model MCS, or selectively only the cells, leaving the ECM unstrained (see sketch in Figure 2a). Dextran molecules with small hydrodynamic radii (r < 6 nm, MW < 70 kDa) penetrate the ECM, without entering the cells ^24^ and thus build up a hydrostatic pressure across their membranes that selectively compresses them. Instead, a global compression is obtained by exposing MCS to an osmotic stress exerted by big Dextran molecules (r > 15 nm, MW > 200 kDa) that are excluded from both cells and ECM (Figure 2b-left) ^14,15^.

**Figure 2:**
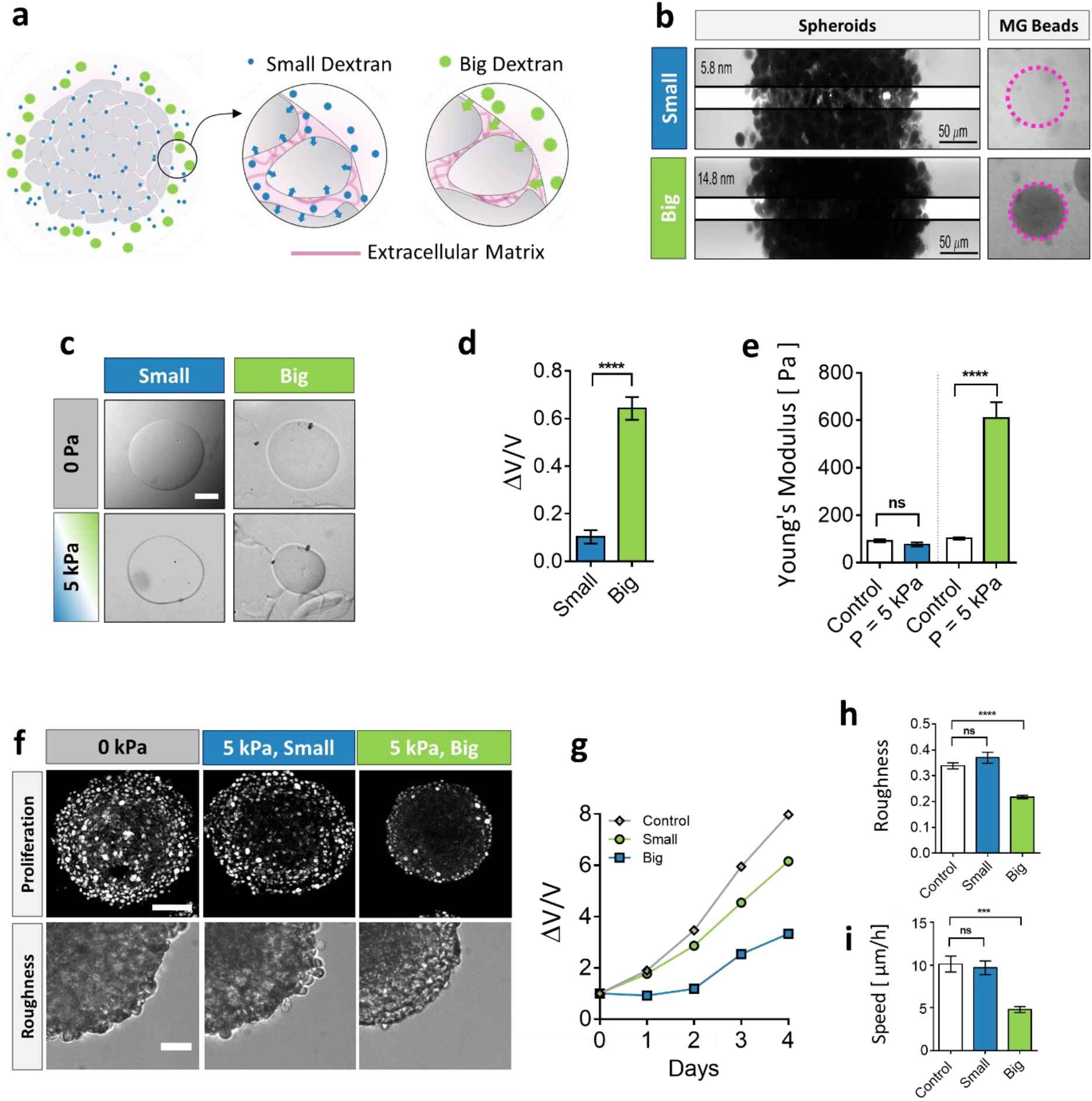
Extracellular Matrix selective compression in MCS. (**a**) Selective compression method. Small Dextran molecules (blue) with MW < 70 kDa and a hydrodynamic radius smaller than 6 nm penetrate the ECM and exert a selective compression on the cells, but have no effect on the ECM. Conversely, big Dextran molecules (green) with MW > 200 kDa (R>15 nm) exert a global compressive stress on the whole MCS. (**b**) Left: Confocal sections of MCS dipped in a solution containing either small or big fluorescent Dextran molecules. The section shows that ECM is selectively permeable to small Dextran, with a cutoff radius of the order of 10 nm. Right: MG beads are also permeable to small Dextran, with a cutoff radius similar to that of ECM. (**c**) Compression of MG beads under 5 kPa, triggered by small and large Dextran molecules. (d) Beads lose 60% of their initial volume when compressed using big Dextran, and only 10% with small Dextran. Error bars represent the standard deviation of the mean. N=10. (**e**) AFM measurement of MG stiffness indicates that, under compression, MG stiffens by 6-fold. N=9 per each condition. (**f**) Above: Proliferating cells revealed by immunostaining of KI67 with no pressure, under overall compression of 5 kPa and under selective compression of the cells and the consequence on MCS growth speed (**panel g**), N > 36 MCS per each condition. Three independent experiments. Below: Effect of compression on the MCS roughness, defined as *Perimeter*/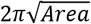and quantification in panel (**h**), N > 36 MCS per each condition. Three independent experiments. (**i**) Cell migration speed (see SI) within MCS also significantly depends on ECM compression. N= 5 independent experiments per condition. All results are displayed as bar graphs ± SD. Biological experiments were repeated at least three times.

First, we assessed the long-term consequences of such global versus selective osmotic compression using three parameters: cellular proliferation, cellular motility and MCS roughness. We observed that whereas cells in control MCS (0 kPa) present a rather uniform proliferation pattern, a global compression of MCS (big Dextran) stops cell division in the core and alters the overall MCS growth (Figure 2f), as previously reported ^9,10,25^. Such altered cellular proliferation under 5 kPa resulted in a three times slower MCS growth as compared to control (Figure 2g). In parallel, cell motility within the MCS at 5 kPa (big Dextran) decreased by 50% (Figure 2i and SI-2). We also observed that global compression induces MCS smoothening (Figure 2f – bottom and Figure 2h), which we assume is related to an increased cell cohesion. Strikingly, all three parameters were not altered when MCS were exposed to an equivalent pressure (small Dextran) applied selectively to cells residing in a native ECM, suggesting the major role of ECM in transducing changes in mechanical state of the environment to cells. However, the ECM properties under compression are difficult to characterize inside the MCS directly.

To probe the rheological properties of the ECM upon osmotic compression, we used Matrigel (MG), an ECM proxy secreted by EHS mouse sarcoma ^26^ Consistently with native ECM, large Dextran molecules were also excluded from microbeads made of MG (Figure 2b-right and SI-1) suggesting an equivalent effective permeability. Upon 5kPa compression, MG microbeads shrunk by approximately 64±5% (Figure 2c-d), which indicates that even small mechanical perturbations *in vivo* are capable to impose important changes to the residing ECM. As measured by atomic force microscopy (AFM), the Young’s modulus of compressed MG gels is significantly larger (610±70 Pa) as compared to control (103±4 Pa), suggesting a strain stiffening behavior of MG (Figure 2e and SI-3). We observed a significantly smaller compression (10±4%) with small Dextran, which enters the MG^27^. Using this experimental system, we applied compression to individual cells seeded in MG to reproduce a 3D-like environment with no neighbors and confirm our results obtained for whole MCS in order to rule out collective effects stemming from potential cell-cell contacts. Few days after insemination in MG, the control cells (0 kPa) presented long protrusions, reminiscent of the typical morphology of invasive malignant cells. A global compression of MG with large osmolites resulted in strikingly different non-protrusive morphotype, while a selective compression of the same magnitude applied via small osmolites was insufficient to alter the morphotype (Figure 3a and 3b). Our results therefore suggest that the ECM plays the role of a mechanical proxy through which the global MCS compression impacts the cells fate.

**Figure 3:**
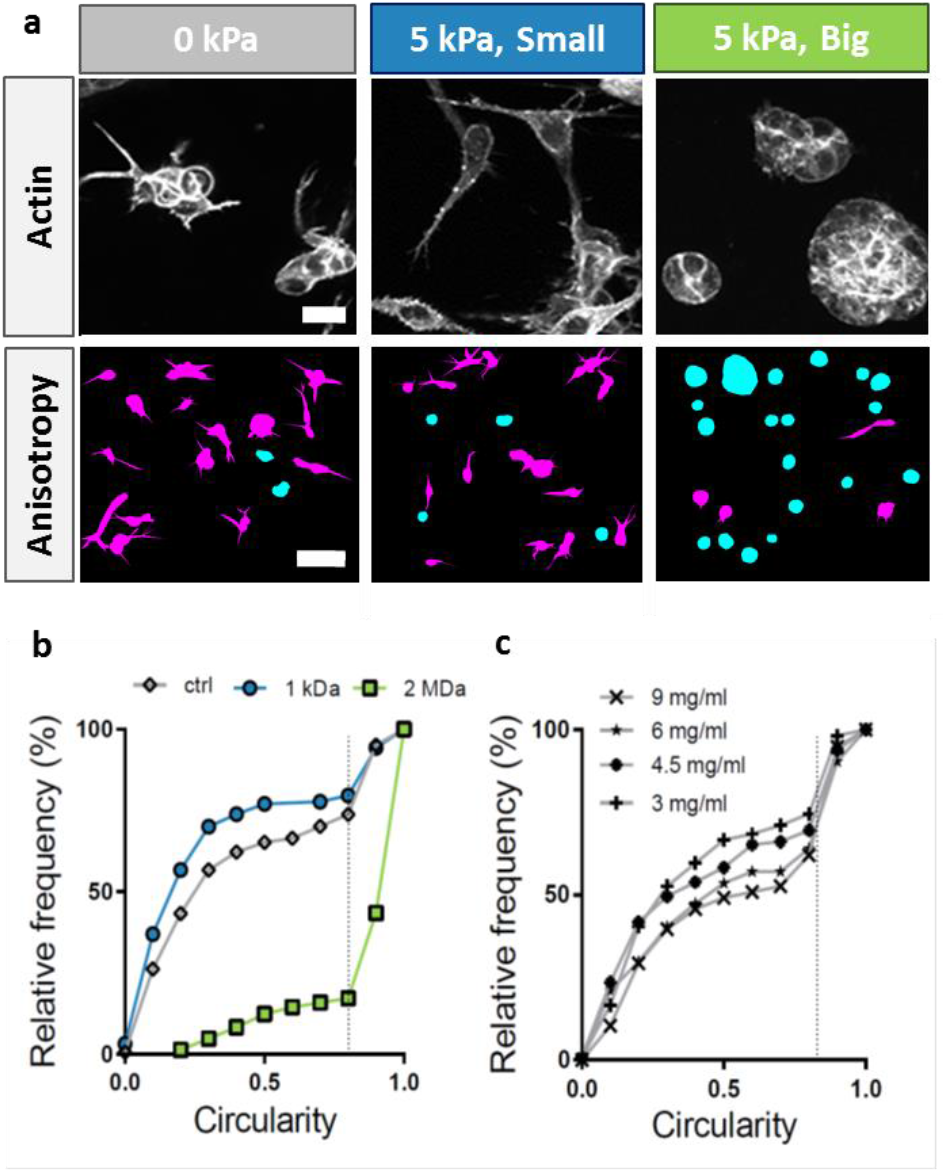
Compression of individual cells in MG. (**a**) MG compression promotes the growth of sphere-like clusters, while cells form stretched structures without pressure, or with a selective pressure but no MG compression. Above: Maximal projection of 50 μm Z-stack, actin staining; Below: analysis of shape anisotropy as a measure of circularity (C); high protrusive phenotype in magenta (C≤0.8) and cluster-like phenotype in cyan (C> 0.8) (**b**) Cumulative distribution of clusters anisotropy in control and under global and selective compression. (**c**) MG initial concentration has little effect on single cell/cluster anisotropy.

To understand the mechanism behind such mecano-sensitivity, in line with experimental observations shown on Figure 1 and Figure 2, we modelled the short time scale response (i.e. associated to water percolation) of the MCS to a global osmotic compression using a composite description, where cells are impermeable and incompressible and the ECM is poroelastic (Figure 4b). A pre-stress, due to the cell division and loss within the MCS, was also taken into account consistently with ^28^ and assumed to be fixed at the timescale of the compression. We obtained (see details in SI-4) the following formulas for the effective volumetric bulk modulus and the hydrodynamic diffusion coefficient associated to water percolation during compression:

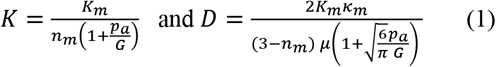

In the above relations, *μ* is the viscosity of the extracellular fluid, *K_m_* is the bulk modulus of the ECM, *κ_m_* the permeability of the ECM, *p_a_* is the active stress in the MCS and *G* its shear modulus. Rigorously, such formulas are derived in a linear theory assuming that *p_a_/G* is small. However, we shall still use them for moderate values of *p_a_/G* ≈ −0.7 as estimated by fitting the theoretically predicted profiles to the experimentally measured local stress and strain ^28,29^ within the MCS (details in see SI-4.3). As we measure by AFM a shear modulus of *G* = 280±10 Pa (Figure d and SI-3.2), we thus estimate an active pressure of the order of *p_a_* = −200 Pa. Both *K_m_* =1.2 ± 0.2 kPa (Figure c and SI-4.4) and *D_m_* = 30 ± 3 *μ*m^2^/s (Figure e and SI-4.4) were directly obtained from the measurement of the compressibility and relaxation time of MG beads under osmotic pressure. Together with the measured value of *n_m_*, formulas (1) have no adjustable parameter and predict a value of *K* of few tens of kPa and *D* of few tens of *μ*m^2^/s. Such values are consistent with direct measurements of *K* = 29±3kPa (Figure c and SI-4.3) and *D* = 44±4 μm^2^/s (Figure e and SI-4.3) based on global compression experiments confirming the model applicability. In this framework, the ECM plays the role of a sensor transmitting the small compressive stress to the cells in the aggregate. Indeed, a global compression increases the hydrostatic pressure in the inter-cellular space, draining the water out and straining the ECM (Figure 4f), which in turn mechanically compresses the cells in a permanent way as such stress would be relaxed only with complete remodeling of the ECM. In contrast, cells rapidly (1-10min) offset a small osmotic stimulus −5 kPa (this corresponds to 1 mM NaCl, which is a very small variation compared the physiological values of concentration of [Na],[Cl]≈150 mM in the extra-cellular space) through a regulatory response, which activates ion pumps ^30^ to equilibrate the internal/external osmolarity unbalance. As a consequence, an omosotic shock selectively compressing the cells has a fundamentally different biological signature on MCS than a global osmotic shock, which exposes the cells to a permanent mechanical pressure.

**Figure 4:**
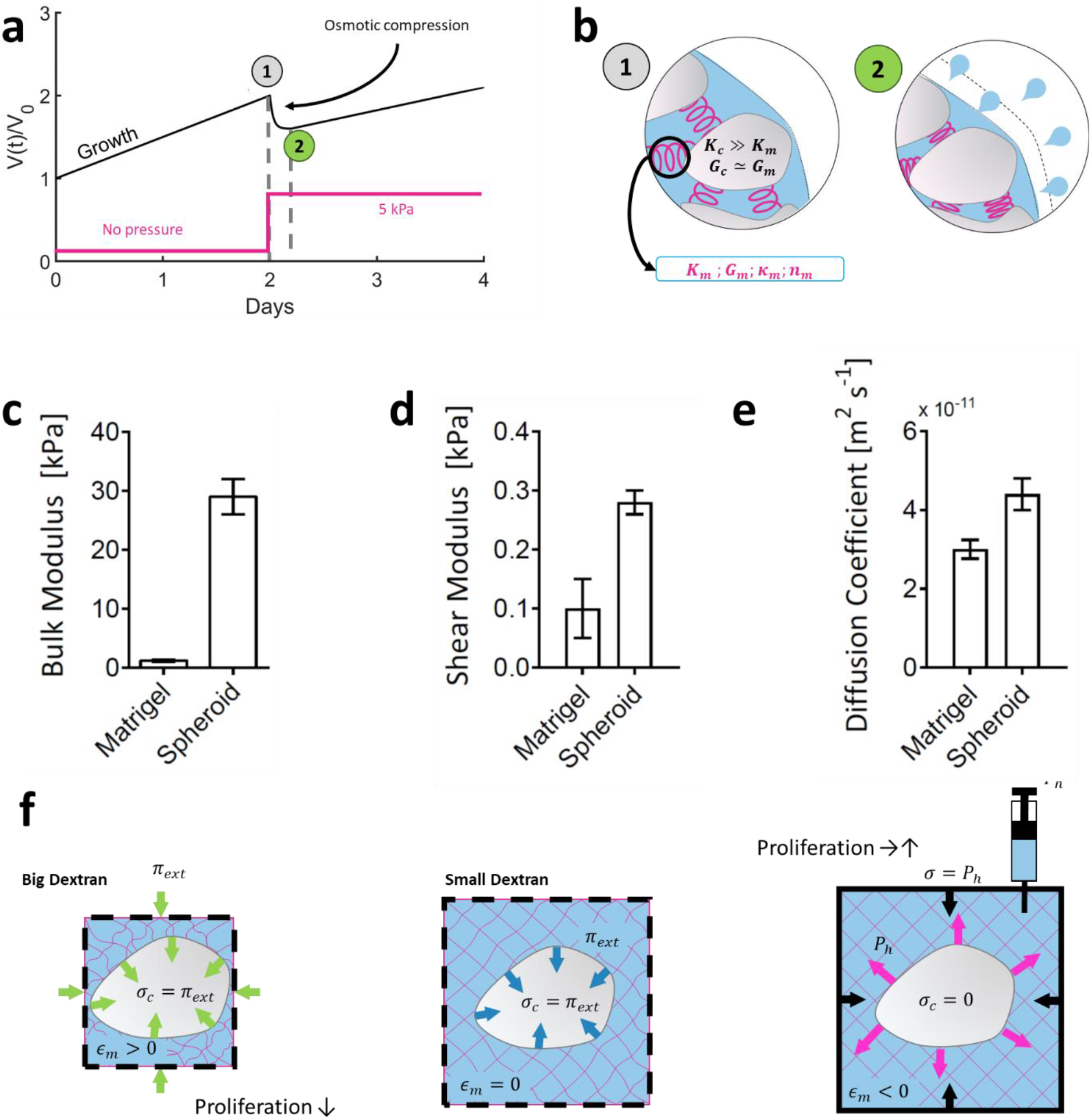
Mechanical response of MCS and MG to mechanical stresses. (**a**) Schematic description of the experiment. The osmotic compression (magenta) is applied after 2 days of unconstrained growth (black curve). The MCS volume decrease typically lasts after few minutes, between marks (1) and (2). (**b**) The MCS is modeled as a composite material made of incompressible cells (bulk modulus K_c_ ~ 1 MPa) surrounded by a porous ECM. ECM is characterized by its bulk and shear moduli (K_m_ and G_m_), its volume fraction n_m_ and the porosity κ_m_. Marks (1) and (2) correspond respectively to the conformations before and after compression (see panel a) (**c**) The bulk modulus of MG and MCS (K_m_ = 1.2±0.2 kPa and K = 29±3 kPa) and (**d**) the corresponding shear moduli (G_m_ = 100±50 Pa and G = 280±10 Pa) measured by AFM as described in SI. (**e**) Diffusion coefficient of water through a bead of pure MG (D_m_ = (3.0±0.2)·10^−11^ m^2^/s) and through the composite MCS (D = (4.4±0.4)·10^−11^ m^2^/s), under compression. (**f**) Propagation of three different stresses in a composite poroelastic MCS. Left: Big Dextran exert the stress on cells through ECM compression (ε_m_ > 0), which cannot be actively screened by cells over a short timescale. Proliferation decreases significantly. Middle: Small Dextran molecules exert the stress directly on the cells, but with no ECM deformation (ε_m_ = 0), and has little impact on proliferation. Indeed the stress is suppressed over a long timescale by an active pumping of ions inside the cell to re-equilibrate the osmotic pressure. Right: ECM hydration observed in in-vivo tumors induces ECM distension (ε_m_ < 0) and an overall MCS volume increase which promotes tumor growth.

The model also indicates that water percolation through the MCS pores effectively follows a diffusive process. Thus, the timescale at which the incompressible/compressible transition occurs scales quadratically with the typical size. In vision of that, tissues are incompressible because of the presence of water but, with the measured value of *D*, an individual cell (typically 20 μm) ‘feels’ an incompressible environment only for few minutes if it swells or moves. As a result, the slow processes of cell division and motion take place in an effectively compressible environment, but where some residual stress stemming from the ECM deformation remains until the ECM is completely remodeled.

## Discussion

Several hypotheses can be put forward as to how cells transduce such residual stress into a biological response. First, MG compression is associated with its stiffening, as the Young’s modulus increases 6-folds (Figure 2e). This would be coherent with the evidence that the substrate stiffness influences the cellular fate^16^. Compression also results in a reduction of MG porosity by ~25% (Figure 2c and SI-1), with the consequent reduction of diffusion of oxygen, nutrients and metabolites as well as insoluble cues present in the ECM (e.g. GPB, cRGD) ^31^. However, for single cells seeded in MG subjected to a global compression, the invasive morphotype was not rescued with MG of higher densities corresponding to the compressed state (Figure 3c), suggesting that the presence of the stress in the external environment directly induces the morphotype change rather than its stiffness or porosity.

Second, ECM compression may restore cell-cell contact inhibition of proliferation^8^ and locomotion^32^ through maturation of cell-cell adhesion. Indeed, as shown by phase contrast images in Figure 2f and quantified in Figure 2i, the whole MCS compression smoothened its surface, which we interpret as an increase of cell-cell adhesion. Conversely smoothing did not occur with small osmolites. The fact that mechanical compression arrests cell motility is also in agreement with a simple model of the cell cytoskeleton self-organization^33^.

Interestingly, a recent study points to the role of the ECM swelling in tumor growth^13^. Because the ECM of tumors contains negatively charged macromolecules, counter-ions permeate the MCS to establish electroneutrality and create an osmotic pressure difference between the MCS inside and outside. This pressure difference is equilibrated by an interstitial hydrostatic pore pressure within the MCS that swells the ECM. It is then conceivable that such deformation, opposite to the ECM compression that we measure during osmotic compression, sustains the MCS growth. Such an effect can be included in our theoretical model by accounting for the ionic species in solution in the MCS pores^34^, however we neglected it given that high (200 mM NaCl) modifications of counter ions concentrations are necessary to record a non-negligible pore pressure increase while we imposed much smaller osmotic perturbations (5 kPa corresponds to 1 mM NaCl). The viability of cells in such osmotic environment would need to be carefully scrutinized.

To conclude, our experiments are compatible with the model proposed here, where ECM mechanical properties along with volume exclusion due to incompressible cells determines the MCS final volume loss and its dynamics upon a gentle global compression. In this framework, the ECM is a sensor through which the global MCS compression is transformed into a permanent biochemical signal impacting proliferation (Figure 4a). While we have shown that the MCS rheology and, in particular, the state of compression of the ECM affect cellular fate within the MCS, cellular turnover will in turn modify the active pre-stress leading to an emergent hydrodynamic diffusion and mechanical stress within the MCS in the long timescale. We therefore anticipate that mechanical theories aiming at capturing such state ^35^ should further account for the presence of the ECM.

## Materials and Methods

### Cell culture, MCSs formation, and growth under mechanical stress

CT26 (mouse colon adenocarcinoma cells, ATCC CRL-2638; American Type Culture Collection were cultured under 37°C, 5% CO_2_ in DMEM supplemented with 10% calf serum and 1% antibiotic/antimycotic (culture medium).

Spheroid were prepared on agarose cushion in 96 well plates at the concentration of 500 cell/well and centrifuged initially for 5 minutes at 800rpm to accelerate aggregation. After 2 days, Dextran (molecular mass 1, 10, 40, 70, 100, 200, 500 and 2000 kDa; Sigma-Aldrich, St. Louis, MO) was added to the culture medium to exert mechanical stress, as previously described ^15^, at a concentration of 55 g/L to exert 5 kPa.

To follow spheroid growth over the time, phase contrast images were taken daily starting from “day 0” before addition of dextrans and after 30 minutes. Spheroid were kept under constant pressure over observation period. Images are analysed using Imagej plugin (doi.org/10.1371/iournaLpone.0103817). Each experiment was repeated 3 times, with 32 individual spheroids per condition.

### Cryosectioning and Immunostaining (fibronectin and KI67)

Spheroids were fixed with 5% formalin (Sigma Aldrich, HT501128) in PBS for 30 min and washed once with PBS. For cryopreservation spheroids were exposed to sucrose at 10% (w/v) for 1 hour, 20% (w/v) for 1 hour and 30% (w/v) overnight at 4°C. Subsequently spheroids were transferred to a plastic reservoir and covered with Tisse TEK OCT (Sakura) in an isopropanol/dry ice bath. Solidified samples were brought to the cryotome (Leica CM3000) and sectioned into 15μm slices. Cut layers were deposited onto poly-L-lysine coated glass slides (Sigma) and the region of interest was delineated with DAKO pen. Samples were stored at −20°C prior immunolabelling. For fibronectin and Ki67 staining samples were permeabilized with Triton X 0.5% in TBS (Sigma T8787) for 15 minutes at RT. Nonspecific sites were blocked with 3% BSA (Bovine serum Albumin) for 1 hour. Then, samples were incubated with first antibody (Fibronectin, Sigma F7387, 1/200 and Ki67; Millipore ab9260, 1/500) overnight at 4°C. Subsequently samples were thoroughly washed with TBS three times, for 15 minutes each. A second fluorescent antibody (goat anti-mouse Cy3, Invitrogen; 1/1000) was incubated for 40 minutes along with phalloidin (1/500, Alexa Fluor 488, Thermo Fisher Scientific). After extensive washing with TBS (four washes of 15 minutes) glass cover slides were mounted on the glass slides with a Progold mounting medium overnight (Life Technologies P36965) and stored at 4°C before imaging.

### Matrigel beads preparation

Matrigel beads were prepared using vortex method. Oil phase of HFE-7500/PFPE-PEG (1.5 %w/v) was cooled down to 4°C. For 400 μL of oil, 100μL of Matrigel was added. Solution was vortexed at full speed for 20 seconds and subsequently kept at 37’C for 20 minutes for polymerization. Beads were transferred to PBS phase by washing out the surfactant phase with pure HFE-7500 oil.

### Matrigel beads compression

To compress polimerized Matrigel beads, PBS was enriched with either 2MDa or 1kDa dextran at the concentration to exert a pressure of 5 kPa. Images were taken just before, and then 45 minutes after, dextran was added. Volume decrease was measured for 10 different beads.

### Matrigel porosity

18 hours before observation, fluorescent dextrans were added to PBS containing Matrigel beads (in control or compressed as previously described) to the final concentration of 500 nM for 500kDa dextran, and 5 μM for 40kDa and 70kDa dextran. Images were taken 30μm above the glass surface by the border of the Matrigel bead. Infusion efficiency was quantified by measuring the normalized Intensity (ImageJ) in at least 9 small regions within Matrigel (I_gel_) and outside in the solution (I_so1_). At least 8 images were taken per condition.

### Infusion of dextrans into Matrigel beads

18 hours before observation, Fluorescent dextrans were added to PBS containing Matrigel beads (in control or compressed as previously described) to the final concentration of 500 nM for 500kDa dextran, and 5 μM for 40kDa and 70kDa dextran.

### Statistical analysis

Student’s t-test (unpaired, two tailed, equal variances) was used to calculate statistical significance as appropriate by using PRISM version 7 (graphpad Software). Statistical significance was given by *, P<0.05; **, P<0.01; ***, P<0.001; ****, P<0.0001.

## Supporting information

## Acknowledgments

We warmly thank J. Prost and F. Jülicher for drawing our attention to the potential impact of the poroelasticity on the rheology of multicellular aggregates.

## Author contributions

M.E.D. and G.C. designed the experiments; M.E.D., B.B., M.L., C.V., M.G. and G.C. performed the experiments; M.E.D., P.R. and G.C. analyzed the data; P.R. and J.-F. J. developed the theoretical model; M.E.D., P.R. and G.C. wrote the manuscript.

