## Supplementary material for "Extracellular Matrix acts as pressure detector in biological tissues"

December 5, 2018

#### 1 Extra-cellular Matrix (ECM)

##### 1.1 ECM volume fraction

To evaluate the volume fraction of ECM in multi-cellular spheroids (MCS), we supplement the culture medium with sulforhodamine-B, a hydrophilic fluorophore that stains the extracellular space without penetrating the cells. From confocal sections of MCS (Fig. 1 a) we determine the thickness of the thin layer between two adjacent cells. By fitting the intensity profile to a Gaussian distribution (Fig. 1 c), and taking into account that the instrumental function (resolution 270nm) broadens the profile, we estimate the extracellular layer to 0.9±0.1 $\mu$m (histogram in Fig. 1 d; N=132). With an average cell diameter of 20 $\mu$m, we evaluate that the fraction of extracellular space is approximately $n_m = V_m/V_0 = 14 \pm 5\%$. An alternative method to evaluate the fraction $n_m$ is to threshold the fluorescence intensity measured in a confocal section (Fig. 1 a). From the number of white pixels in the images after thresholding (Fig. 1 b), we estimate that the fraction of extracellular space goes to $n_m = 14 \pm 4\%$. The experimental uncertainty is due to the empiric choice of parameters used in the threshold process. Fig. 1 e compares the results obtained with the two methods. The two methods give similar results and exhibit a non-negligible volume fraction of ECM in MCS.

##### 1.2 Exclusion size of Matrigel (MG)

Electron microscopy observations show that pore sizes are extremely heterogeneous in hydrogels. Thus, the typical pore size of MG is difficult to evaluate. However, we can empirically define an exclusion-size, above which globular molecules do not penetrate the gel. To evaluate this exclusion-size, we dip MG beads in a solution containing fluorescent tracers with different radii. Depending on their size, those tracers either enter the MG or not. In practice, we use fluorescently labelled Dextran polymers with different molecular weights (40, 70 and 500 kDa), corresponding to Stokes' radii ($R_S$) of 4.4 nm, 5.8 nm and 14.8 nm respectively. At very low concentration ($< 5\mu$M), the fluorescent Dextran does not substantially contribute to the osmotic pressure. To avoid confusion between Dextran molecules used to apply the osmotic stress (large molecular weight, high concentration, non-fluorescent) and Dextran tracers (small molecules, low concentration and fluorescently labelled), we will refer to the latter simply as *tracers*. Fig. 2 shows that small tracers with $R_S < 5.8$ nm fully permeate MG beads, as we observe the same level of fluorescence both inside the MG beads and in the surrounding solution. Conversely, beads dipped in a solution containing large tracers ($R_S = 14.8$ nm) appear darker than the surrounding medium. Large tracers are excluded from the MG. Our results indicate that the MG exclusion-size is in the range between 6 and 14 nm.

##### 1.3 Exclusion size of ECM in MCS

We observe a similar behavior in MCS dipped in a culture medium supplemented with the same tracers. As shown in Fig. 3 A, tracers with $R_S = 4.4$ nm and $R_S = 5.8$ nm permeate the

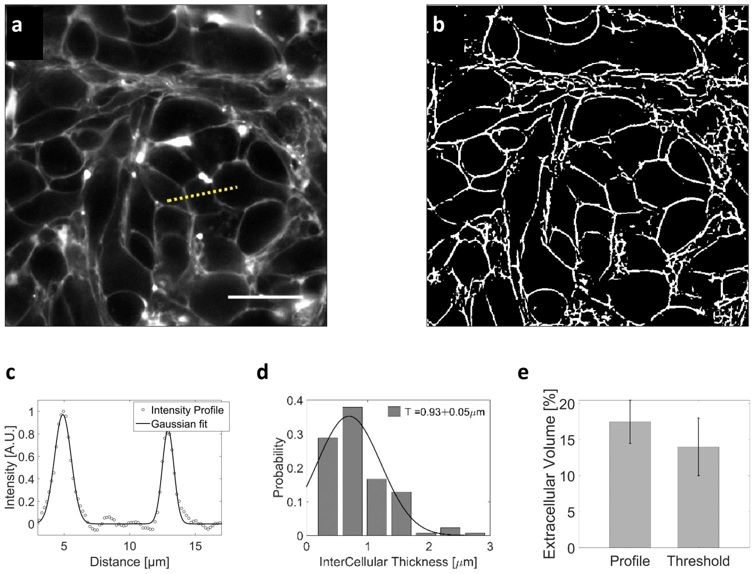


Figure 1: Volume fraction estimation. (a) Thickness method: confocal section of an uncompressed MCS, the extracellular space of which is filled with sulforhodamine-B, a hydrophilic fluorophore that does not enter the cells. The intensity profile across two extracellular layers (panel c). The width of intercellular space is computed by fitting the intensity to a gaussian profile and displayed on panel (d). (b) Threshold methods: A threshold is applied to Images (a) and (b) respectively to evaluate the ratio between extracellular (white) and intracellular (black) space. Depending on the choice of the threshold, the ratio of extracellular space varies by ±5%. (e) Summary of the results obtained using the two methods. Whereas the experimental error is large, the observations show that the extracellular space represents about 15% of the total MCS volume.

extracellular space of the MCS but not those larger than 14.8 nm. In order to quantify the relative amount of tracers inside the MCS, we compare the average fluorescence measured inside the MCS $\langle I_{In} \rangle$ and in the surrounding solution $\langle I_{Out} \rangle$. Fig. 3 B and Fig. 3 C report the relative intensities $\langle I_{In} \rangle / \langle I_{Out} \rangle$ , obtained respectively at an external osmotic pressure $\Pi_e = 0$ Pa (180 MCS) and at $\Pi_e = 5$ kPa (43 MCS). In both cases, the fluorescence level significantly lowers with large tracers. In terms of pore size, MG is thus a good proxy of ECM, as both have an exclusion size of about 10 nm.

#### 2 Pressure and Motile Activity inside MCS

The compressional state of the MCS affects the motility of constituent cells [8, 1]. Recently, we developed a method to measure this motility without using confocal microscopy, which is limited in terms of sample thickness and observation time. In our setup [5] (Fig. 4 a), the MCS is observed by phase contrast (Fig. 4 c) and is simultaneously illuminated with an infrared laser (850 nm). The MCS produces the single-scattering dynamic interference pattern shown in Fig. 4 b (Dynamic Light Scattering). From the temporal fluctuations of this pattern (Fig. 4 d Fig. 4 e), one computes the average velocity of moving cells inside the MCS. It has to be noticed that Dynamic Light Scattering provides information on the 3D motility, and not only on the 2D motion as previously measured by Alessandri et al. at the surface of MCS [1]. With this method, we measure the average speed in the three cases of interest: without pressure, when a pressure is selectively applied on the cells, but not on the ECM (small Dextran), and when

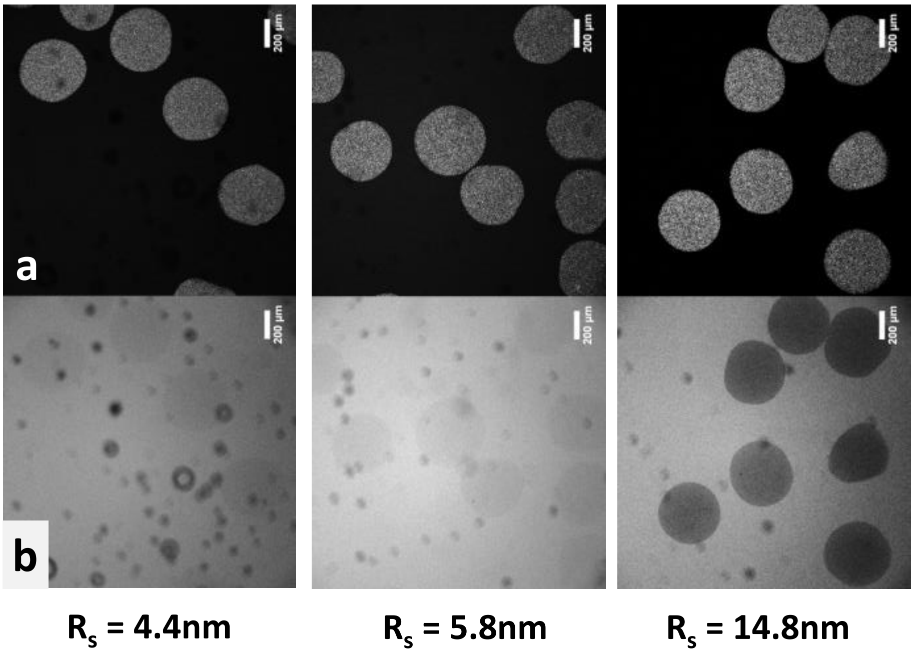


Figure 2: a) MG beads observed by epifluorescence. b) MG beads dipped in a solution containing fluorescent tracers of different sizes. Small tracers ($R_S < 5.8$ nm) penetrate the MG beads, while larger tracers ($R_S$=14.8 nm) are excluded. Scale bars = 200$\mu$m.

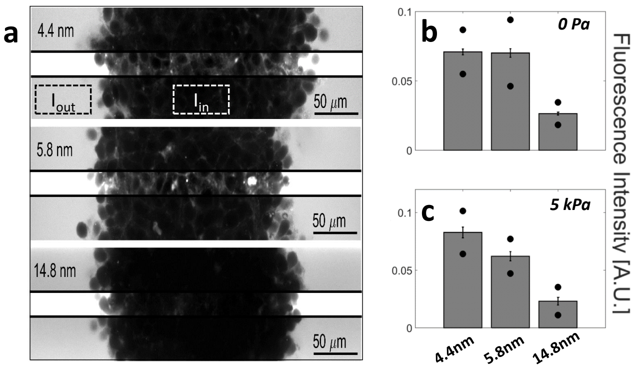


Figure 3: Exclusion size of the ECM in MCS. a) Confocal sections of three MCS dipped in culture media supplemented with Dextran of increasing molecular weights. In order to quantify the total amount of Dextran permeating the MCS, the mean fluorescent intensity measured inside the MCS $\langle I_{In} \rangle$ is normalized by the mean intensity measured in solution $\langle I_{Out} \rangle$. To avoid saturation of $\langle I_{Out} \rangle$, photomultiplier gain is kept low. This reduces the visibility of extracellular space inside the MCS. In the middle stripe of each image, the brightness is increased of the same amount to make the fluorescence of Dextran visible in the extracellular space. b) Relative intensity $\langle I_{In} \rangle / \langle I_{Out} \rangle$ for different Dextran sizes and without osmotic pressure. Black circles and error bars represent respectively the standard deviation and the standard error of the mean, computed over more than 58 MCS per condition. c) Relative intensity with an additional osmotic pressure of 5 kPa, averaged over 16 (for 4.4 nm), 14 (for 5.8 nm) and 13 (for 14.8 nm) MCS.

the pressure is applied to the whole MCS (big Dextran). The results are shown in Fig. 4 f and Fig. 4 g: whereas the average speed is comparable (10±1 $\mu$m/h) in the first two cases (magenta and cyan), it is reduced by a factor of two when the compression is exerted on the entire MCS (4.6±0.3 $\mu$m/h, blue). This result confirms the role of the ECM compression, on how fast cells migrate between the other cells inside a MCS.

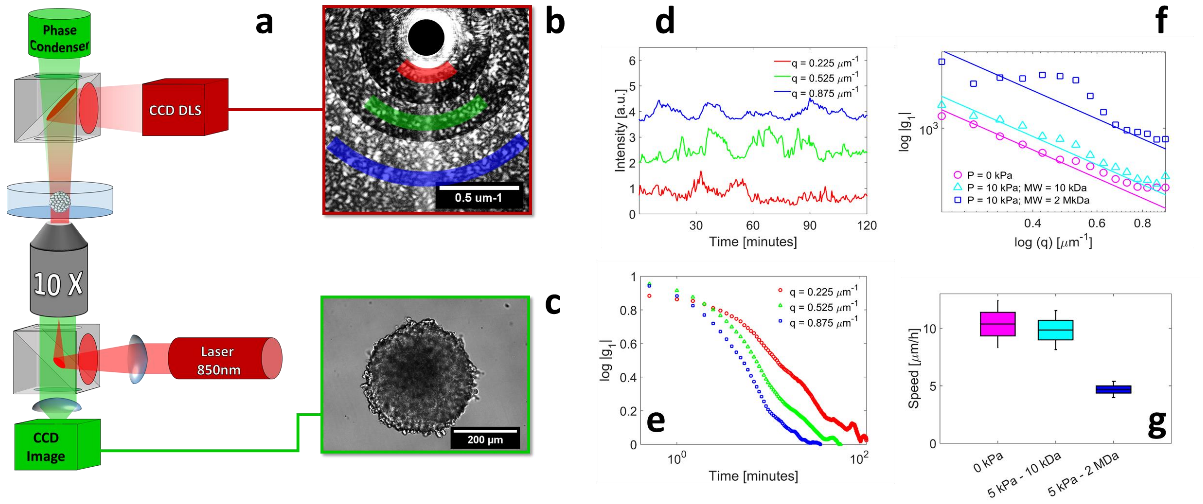


Figure 4: . Motile activity measured by Dynamic Light Scattering (DLS) under different pressure conditions. a) The experimental setup combines two counter-propagating optical pathways, with two different wavelengths. This allows us to observe the MCS simultaneously by DLS ($\lambda_{DLS}$ = 850nm, dark red in the sketch) and by phase contrast ($\lambda_{phase}$ = 530nm, green). The DLS signal and phase contrast image are illustrated respectively in panels (b) and (c). DLS signals are acquired at different scattering vectors **q** and averaged over rings of equal q=|**q**| (colored sectors in panel (b) ). d) Time evolution of diffraction intensity at three different **q**; colors correspond to that of sectors in panel (b). e) Intensity-Intensity autocorrelation functions for the three different scattering vectors. In the single-scattering regime, the intensity signal decorrelates in a typical timescale $\tau = 1/qv^0$, where $v^0$ is the mean cell velocity inside the MCS. f-g) respectively the decorrelation time as a function of q and the resulting mean cell velocity, measured in three different conditions: with no pressure, $v^0$ = 10±1 $\mu$m/h (magenta), with 5 kPa exerted by small Dextran, $v^0$ = 9.8±0.8 $\mu$m/h (cyan), and with $\Pi_e$ = 5 kPa exerted by large Dextran, $v^0$ = 4.6±0.3 $\mu$m/h (blue). Box sizes correspond to the standard error of the mean (N = 5 MCS) and the error bars to the standard deviation.

### 3 Young and Shear moduli

## 3.1 MG

To measure the evolution of MG Young modulus with the compressive stress, we prepare flat sheets of MG, by positioning a drop of 40$\mu$L of MG onto a surface of a petridish and by covering it with a hydrophobic plastic film of surface area $\simeq 1cm^2$. The petridish is then kept at 37°C for 20 minutes to favour MG polymerization and, eventually, the hydrophobic film is carefully removed. MG gels are prepared at least 1.5 hour before the experiment and remain in PBS to avoid gel swelling during AFM measurements (JPK Nanowizard II mounted on inverted microscope C. Zeiss, Observer D1). For compression, 1kDa and 2MDa Dextrans at corresponding concentrations are added directly onto gels. AFM measurements are performed 20 minutes after. We use the same AFM (Bruker, Cantilever C, $k_C = 0.013$ N/m) to measure Young modulus. The approach speed is set to 1 $\mu$m/s. Experiments are repeated on three different gels per each condition. Errors corresponding to Data are using the JPK analysis software and built in Hertz model. In Fig. 5, error bars represent the standard deviation. T-test is used for statistical analysis.

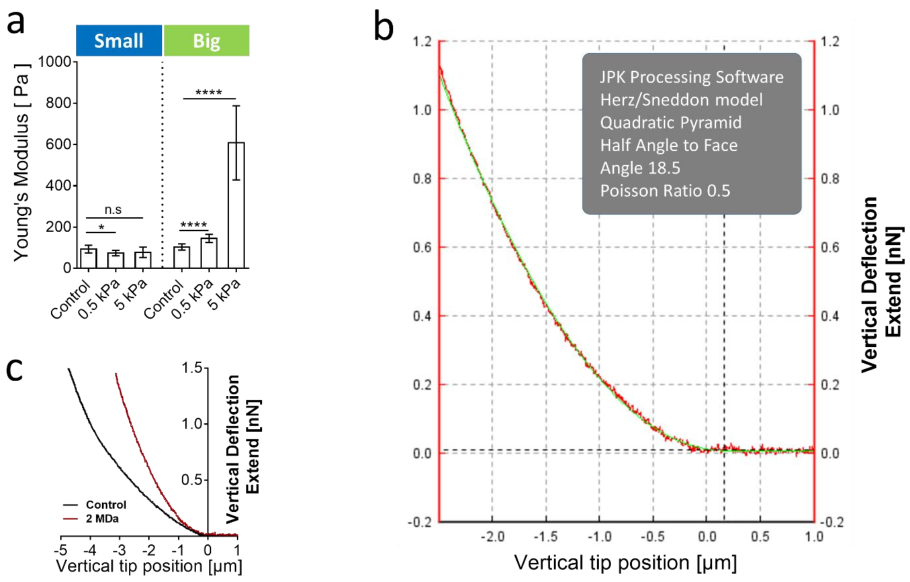


Figure 5: (a) Young modulus for MG beads under increasing osmotic stresses. The Young modulus increases with stress, when the compression is occasioned by big Dextran molecules (right). Conversely, small Dextran molecules do not affect the Young modulus of MG. (b-c) raw data from AFM measurements (indentation-force curves). The curve displays the indentation force $F$ as a function of the vertical tip position.

#### 3.2 MCS

To determine the shear modulus of the MCS $G_s$, we use a the same AFM as for MG. The tipless nominal cantilever stiffness is k = 30 N/m (Nanosensors, TL-NCL-10) and the MCS radii are in the range 100-130 $\mu$m (Fig. 6 a). The indentation speed varies between 20 and 40 $\mu$m/s. At long timescale, the MCS relaxes with a typical time $\tau = 59 \pm 3s$ (standard error of the mean, N = 4 MCS). This value corresponds to the poroelastic timescale, due to water percolation. To measure the undrained Young and the Shear moduli, we analyze the initial part of the Indentation-Force curve. At this timescale ($<$ 500 ms), much shorter than the poroelastic transition, the MCS can be considered as undrained. At small indentation ($<$ 10 $\mu$m), the curve fits well to Hertz's model (Fig. 6 b) for a value of the undrained Young modulus $E^u = 840 \pm 20$ Pa and, consequently, $G = 280 \pm 10$ Pa (using an infinite value for the undrained bulk mudulus $K_u$, defined in the following section). This value is consistent with that found by Guevorkian et al. [12] on similar MCS.

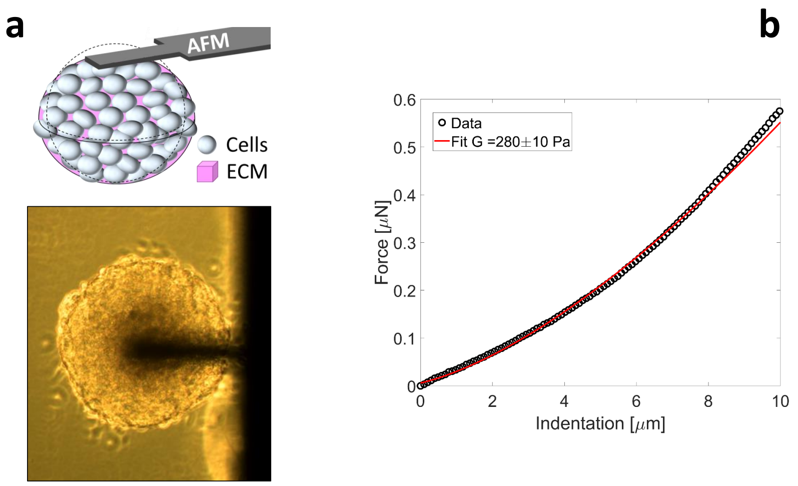


Figure 6: (a) Sketch and image of a MCS in contact with the AFM tip. (b) Force response of a MCS indented at constant speed (20 $\mu$m/s) and fitted using Hertz's model. The best fit is obtained for a shear modulus $E^u = 840 \pm 20$ Pa.

### 4 Model for the deformation of a pre-stress MCS under osmotic pressure

Poroelasticity is a theory describing the flow of a liquid in a solid deformable porous medium. The premise of the theory dates back to Gibbs and was then pioneered by M.A. Biot [4]. A justification of the theory from a homogenization perspective as well as its thermodynamic foundations can be found in the book of O. Coussy [6].

#### 4.1 The mechanical problem

We consider the active poro-elastic framework formulated in [10] to describe the mechanical response of the MCS subjected to an osmotic shock. Before application of the osmotic shock, the MCS occupies the domain $\Omega$ of center 0 and radius $R_m$. We denote the space coordinate $\underline{X}$ in $\Omega$ and the time $t > 0$. The boundary of $\Omega$ is denoted $\partial\Omega$ and the outward normal at a point $\underline{X}$ of the boundary is $\underline{N}$.

**Kinematics.** Because of external loading, the MCS is deformed from its initial configuration $\Omega$ at time $t = 0$ to a new configuration $\omega$ at time $t$. Subsequently, material points at position $\underline{x} \in \omega$ are mapped from the initial configuration by the transformation $\underline{x} = \phi(\underline{X}, t)$ whose gradient is called the deformation tensor $\mathbb{F}(\underline{X}, t) = \nabla_X \phi$.

To account for the presence of a pre-existing incompatibility leading to residual pre-stress prior to the deformation we adopt the classical framework of the multiplicative decomposition of the deformation [19], $\mathbb{F} = \mathbb{F}_e \mathbb{F}_g$ where $\mathbb{F}_e(\underline{X}, t)$ corresponds to the poro-elastic deformation happening at a short timescale after the osmotic shock while $\mathbb{F}_g(\underline{X})$ is an active deformation which builds up at long timescale and can be considered as fixed at the timescale of the experiment. In this paper, we primarily associate this active deformation to cells turnover (division/apoptosis) but it can also be from a different origin including cell contractility or active adhesion. In such cases however, it is less clear that the timescales related to the poro-elastic compression and the build up of active stress are well separated.

Note that both $\mathbb{F}_e$ and $\mathbb{F}_g$ may be incompatible, i.e. not be the gradient of vector field [11] while $\mathbb{F}$ is necessarily compatible by definition. Within this framework, we now write the fundamental physical balance laws.

**Momentum balance.** The first Piola-Kirchhoff stress tensor $\mathbb{P}(\underline{X}, t)$ satisfies the force balance equation,

$$\operatorname{div}(\mathbb{P}) = 0 \text{ with boundary condition } \mathbb{P}|_{\partial\Omega}\underline{N} = 0. \tag{1}$$

Note that we measure the stress with respect to the hydrostatic pressure in the external fluid, which implies the absence of traction force at the MCS boundary. The balance of torques imposes the symmetry condition on the stress $\mathbb{F}\mathbb{P}^T = \mathbb{P}\mathbb{F}^T$ (see for instance [2]).

**Mass balance.** Assuming that the internal flow of extra-cellular fluid follows a Darcy law, mass conservation of the incompressible fluid can be expressed in the initial configuration (see [3] for details) as

$$\frac{\partial n}{\partial t} - \frac{\kappa}{\mu}\nabla^2 p = 0, \tag{2}$$

where $n(\underline{X}, t)$ is the porosity (dimensionless) of the MCS (i.e. the local proportion of interstitial volume), $\kappa$ is the MCS effective permeability (m$^2$) , $\mu$ (Pa.s) is the extra-cellular fluid viscosity and $p(\underline{X}, t)$ the hydrostatic pressure in the pores. At the timescale of interest, we neglect the exchange of water between cells and extra-cellular medium through aquaporins compared to the water permeation in the MCS pores.

To derive the boundary conditions (B.C.) associated to (2), we conceptually imagine the presence of a very compliant dyalisis bag around the MCS which lets water and ions go through but is impermeable to large macromolecules used to perform the osmotic shock. If the filtration coefficient of the bag is denoted by $L_p$, the water flux inside the bag is $L_p(\triangle\Pi - \triangle P)$ where $\Pi$ and $p$ are the osmotic and hydrostatic pressures and $\triangle$ denotes the difference between the two sides of the dyalis bag [15]. Taking the limit $L_p$ large (infinite permeability of the bag), as the external hydrostatic and internal osmotic pressures vanish, we obtain

$$p|_{\partial\Omega} = -\Pi_e,$$

where $\Pi_e(t)$ is the external osmotic pressure controlled experimentally.

**Constitutive behavior.** For a poro-elastic material, we relate the stress to the elastic strain and the pore pressure in the following way [10]:

$$\mathbb{P} = \det(\mathbb{F}_g)\mathbb{F}_e\mathbb{S}(\mathbb{F}_e)\mathbb{F}_g^{-T} - \alpha\det(\mathbb{F})\mathbb{F}^{-T}p, \tag{3}$$

where $\alpha$ is the (dimensionless) Biot coefficient. We assume that the elastic second order Piola-Kirchhoff stress tensor reads, according to Hooke's law,

$$\mathbb{S}(\mathbb{F}_e) = 2G\mathbb{E}_e + (K_d - 2G/3)\operatorname{tr}(\mathbb{E}_e)\mathbb{I},$$

where $K_d$ (Pa) is the drained bulk modulus of the MCS (i.e. the modulus measured if water can flow out during deformation), $G$ (Pa) its shear modulus and $\mathbb{I}$ the identity tensor. The geometrically non-linear strain measure is the conventional right Cauchy-Green tensor, $\mathbb{E}_e = (\mathbb{F}_e^T\mathbb{F}_e - \mathbb{I})/2$.

A second thermodynamic relation gives the change of porosity to the elastic strain and the pore pressure

$$n - n_0 = \alpha\det(\mathbb{F})\operatorname{tr}(\mathbb{E}_e) + M^{-1}\det(\mathbb{F})p, \tag{4}$$

where $n_0(\underline{X})$ is the porosity in the absence of elastic deformation and pore pressure and $M$ (Pa) is a mechanical modulus.

**Simplifications.** We can re-express the three rheological coefficients $K_d$, $\alpha$ and $M$ as functions of the equivalent moduli $K_d$, $K_u$ and $B$ which are respectively the drained and undrained (i.e. measured if the fluid is trapped in the porous medium) bulk moduli and the Skempton coefficient (see [18] for details):

$$\alpha = B^{-1}\left(1 - \frac{K_d}{K_u}\right) \text{ and } M^{-1} = \frac{K_u - K_d}{B^2 K_u^2}.$$

We assume that $B \simeq 1$ and $K_u \gg K_d$ as it is very difficult to compress the MCS in an undrained situation. Thus we obtain that $\alpha \simeq 1$ and $K_d M^{-1} \ll 1$ and we are left with only four rheological parameters, $K_d$, $G$, $\kappa$ and $\mu$ determining the response of the MCS to the osmotic compression.

Finally, we also suppose that the deformation from the initial configuration in response to the osmotic compression remains small, leading to $\mathbb{F} \simeq \mathbb{I} + \nabla \underline{u}$ where $\underline{u}(\underline{X}, t)$ is the displacement field. Thus at first order, the final problem reads,

$$\begin{array}{c} \operatorname{div}(\mathbb{P}) = 0 \text{ with B. C. } \mathbb{P}|_{\partial\Omega}\underline{N} = 0. \\ \frac{\partial \delta n}{\partial t} - \frac{\kappa}{\mu}\nabla^2 p = 0 \text{ with B. C. } p|_{\partial\Omega} = -\Pi_e \\ \mathbb{P} = \det(\mathbb{F}_g)\mathbb{F}_e\mathbb{S}(\mathbb{F}_e)\mathbb{F}_g^{-T} - p\mathbb{I} \text{ and } n - n_0 = \operatorname{tr}(\mathbb{E}_e) \text{ where } \nabla u = \mathbb{F}_e\mathbb{F}_g - \mathbb{I} \end{array} \tag{5}$$

where $\delta n = n - n_0$ is the change of porosity from the initial configuration and $\mathbb{F}_g$ is an imposed quantity resulting from long time cell turnover while the other quantities have to be determined.

#### 4.2 The passive case

In the absence of active cell turnover in the MCS, $\mathbb{F}_g = \mathbb{I}$ and (5) reduces to a classical linear poro-elastic problem where the Cauchy, first and second Piola-Kirchhoff stress tensors all coincide and the Cauchy-Green strain $\mathbb{E}_e = \epsilon = (\nabla \underline{u} + \nabla \underline{u}^T)/2$ is equal to the infinitesimal strain. In this case, taking the divergence of the mechanical equilibrium combined with the two constitutive relations, we obtain, $\nabla^2 p = (4G/3 + K_d)\nabla^2 \delta n$ and (5) becomes

$$\begin{array}{c} \operatorname{div}(\mathbb{S}(\epsilon)) = \nabla p \text{ with B. C. } \mathbb{S}(\epsilon)|_{\partial\Omega}\underline{N} = p|_{\partial\Omega}\underline{N} \\ \frac{\partial \delta n}{\partial t} - \frac{\kappa(4G/3+K_d)}{\mu}\nabla^2 \delta n = 0 \text{ with B. C. } p|_{\partial\Omega} = -\Pi_e. \end{array} \tag{6}$$

From (6) we deduce two classical results:

1. After the osmotic shock, the water percolates out of the MCS following a diffusion process with a diffusion coefficient [21]

   $$D_{\text{passive}} = \kappa(4G/3 + K_d)/\mu,$$

   until a steady state is reached where the hydrostatic pore pressure equates minus the imposed osmotic pressure throughout the whole MCS.

2. In this final state, the volumetric strain within the MCS is constant $\operatorname{tr}(\epsilon) = -\Pi_e/K_d$. Hence, the relative change of volume of the MCS following the shock is associated to the drained compressibility modulus

   $$K_{\text{passive}} = K_d.$$

In the following, we generalize these results to the active case where the MCS grows.

#### 4.3 The active case

To simplify notations, we denote $\mathbb{F}_a = \mathbb{F}_g^{-1}$. Then, in the small displacement case, we can express the stress as

$$\mathbb{P} = \mathbb{P}_a + \nabla \underline{u}\mathbb{P}_a + \det \mathbb{F}_a^{-1}\mathbb{F}_a\mathbb{S}(\mathbb{F}_a^T \epsilon \mathbb{F}_a)\mathbb{F}_a^T - p\mathbb{I} \tag{7}$$

where $\mathbb{P}_a = \det \mathbb{F}_a^{-1} \mathbb{F}_a \mathbb{S}(\mathbb{E}_a) \mathbb{F}_a^T$ is the active stress in the MCS prior to osmotic compression. Again, it is assumed that this stress does not change in time during the short time of the compression since it is set by cell divisions and apoptosis happening on a longer time scale.

Prior to the compression, the deformation is $\mathbb{F} = \mathbb{I}$ but there already exists a pre-stress $\mathbb{P} = \mathbb{P}_a$ in the MCS. As mechanical equilibrium has to be satisfied, $\mathbb{P}_a$ has to verify

$$\mathrm{div}(\mathbb{P}_a) = 0 \text{ with boundary condition } \mathbb{P}_a|_{\partial\Omega}\underline{N} = 0 \text{ and } \mathbb{P}_a = \mathbb{P}_a^T. \tag{8}$$

As (8) in particular implies that $\mathbb{P}_a$ is symmetric, we can rewrite (7),

$$\mathbb{P} = \mathbb{P}_a + \epsilon\mathbb{P}_a + \det \mathbb{F}_a^{-1} \mathbb{F}_a \mathbb{S}(\mathbb{F}_a^T \epsilon \mathbb{F}_a) \mathbb{F}_a^T - p\mathbb{I}. \tag{9}$$

and (5) can be expressed using the infinitesimal strain only:

$$\begin{gathered} \mathrm{div}(\mathbb{P}) = 0 \text{ with B. C. } \mathbb{P}|_{\partial\Omega}\underline{N} = 0. \\ \frac{\partial \delta n}{\partial t} - \frac{\kappa}{\mu}\nabla^2 p = 0 \text{ with B. C. } p|_{\partial\Omega} = -\Pi_e \\ \mathbb{P} = \mathbb{P}_a + \epsilon\mathbb{P}_a + \det \mathbb{F}_a^{-1} \mathbb{F}_a \mathbb{S}(\mathbb{F}_a^T \epsilon \mathbb{F}_a) \mathbb{F}_a^T - p\mathbb{I} \text{ and } n - n_0 = \mathrm{tr}(\mathbb{E}_a) + \mathrm{tr}(\mathbb{F}_a^T \epsilon \mathbb{F}_a) \end{gathered} \tag{10}$$

where $\mathbb{F}_a(\underline{X})$, has to satisfy the mechanical equilibrium (8) and is imposed from a theory of cell turnover in the MCS at a long time scale such as the one presented in [7]. The (measurable) change of porosity from the initial pre-stressed configuration is then $\delta n = \mathrm{tr}(\mathbb{F}_a^T \epsilon \mathbb{F}_a)$.

**Determination of the active strain.** A simple ansatz of $\mathbb{F}_a$ respecting the symmetry of the problem can be expressed in spherical coordinates

$$\mathbb{F}_a = \begin{pmatrix} a_r(R) & 0 & 0 \\ 0 & a_\theta(R) & 0 \\ 0 & 0 & a_\theta(R) \end{pmatrix},$$

where $R$ in $[0, R_m]$ is the radial coordinate. The cell source term (see [8]) can be written $\rho k$ where $\rho$ is the mass density of cells and $k = d\delta/dt$ is the measured rate of cell production, $\delta(R,t)$ being the number of cells appearing from division ($\delta > 0$) or dying through various processes ($\delta < 0$) per unit time. This variable is related to $\mathbb{F}_a$ by mass balance (see for instance [11]) through the relation $\det \mathbb{F}_a = a_r a_\theta^2 = \exp(-\delta)$. With $\delta$ imposed experimentally, $\mathbb{F}_a$ thus depends on a single scalar variable (say $a_\theta$) which can be fully determined by solving the non-linear ODE problem resulting from (8). In general, this operation cannot be done analytically.

However, we can postulate the spatial form of the cell proliferation

$$\delta(R) = \delta_a \left[ (R/R_c)^\beta - 1 \right], \tag{11}$$

where $R_c$ is the critical radius above which cells start to predominantly divide while they are mostly dying for $R < R_c$, $\delta_a$ is the characteristic magnitude of $\delta$ and $\beta$ dictates the spatial gradients of $\delta$. Next, solving (8) with the ansatz (11) in the simple case (linear ODE) where $\delta_a \ll 1$ and $G \ll K_d$ we obtain,

$$a_\theta(R) = 1 + p_a \frac{3\left(\frac{R_m}{R}\right)^3 \left(1 - \left(\frac{R_c}{R_m}\right)^\beta\right) - \beta\left(\left(\frac{R}{R_c}\right)^\beta - \left(\frac{R_m}{R}\right)^3\right)}{4(\beta+3)G},$$

where we define the active pressure as $p_a = K_d \delta_a$. Importantly, this small perturbation regime strictly holds in the limit

$$p_a \ll G \ll K_d,$$

which is assumed in the rest of the text. While experimental data do suggest that $K_d$ is larger than $G$ by an order of magnitude (See Section. 3), the second limit $p_a \ll G$ is less clear but we still use the ensuing asymptotic formulas. In the following, we also investigate the limit when $R_c \ll R_m$ corresponding to a very small necrotic core as our experiments correspond to relatively 'young' MCS (2-3 days of maturation and radii between $100 - 200\mu$m).

Note that we can make an explicit link of the present approach with the theory presented in [7]. Within such theory, the final residual stress is reached when the MCS volumetric expansion stops:

$$\int_0^{R_m} R^2(\mathrm{e}^{\delta(R)} - 1)dR = 0$$

which implies that $R_c = (3/(3+\beta))^{1/\beta} R_m$. Importantly, we need to impose $\beta > -3$ for all quantities to be integrable. In this case, we obtain

$$\mathbb{P}_a = -p_a \begin{pmatrix} \left(\frac{R}{R_m}\right)^\beta - 1 & 0 & 0 \\ 0 & \left(\frac{R}{R_m}\right)^\beta \left(\frac{\beta}{2}+1\right) - 1 & 0 \\ 0 & 0 & \left(\frac{R}{R_m}\right)^\beta \left(\frac{\beta}{2}+1\right) - 1 \end{pmatrix} \tag{12}$$

which is exactly the same expression as the one found in [7] with the correspondence

$$p_a = \frac{3(\zeta_0 + \zeta_1 P_\mathrm{h})(3(\beta+3)\bar{\eta} - 4\beta\eta\nu)}{(\beta+3)\zeta_1\eta_\mathrm{eff}(2\zeta_1\nu - 3)}.$$

See notations in [7] and we have assumed the surface tension $\gamma = 0$.

As in [8] experimental measurements suggest that $\beta$ is negative, we infer that $\delta_a$ and hence $p_a$ are also negative so that cells divide more closer to the surface than in the bulk as experimentally observed [8].

**Stress and strain in the MCS.** Now that $\mathbb{F}_a$ is determined, solving (10) using a perturbation expansion, we compute the relative loss of volume when an osmotic shock is applied by finding the final radial displacement field (i.e. when the pore pressure reaches its final value $-\Pi_e$ uniformly in the MCS),

$$u^\infty(R) = u_0^\infty(R) + \delta_a u_1^\infty(R) \text{ where } u_0^\infty(R) = -\frac{R\Pi_e}{3K_d} \text{ and } u_1^\infty(R) = \frac{\Pi_e R\left(3\left(\frac{R}{R_c}\right)^\beta - \beta - 3\right)}{3(\beta+3)G}.$$

We can thus obtain the local volumetric strain $\mathrm{tr}\epsilon^\infty$ when water has percolated out of the MCS:

$$\mathrm{tr}\epsilon^\infty = -\frac{\Pi_e}{K_d}\left(1 + \frac{p_a}{G}\left(1 - \left(\frac{R}{R_c}\right)^\beta\right)\right). \tag{13}$$

The polyacrylamide beads that serve as local stress gauges in [9] are internalized within the MCS before the osmotic stress is applied and are permeable to water such that the pore pressure equilibrates between the intercellular space and the beads. They are therefore sensitive to the stress field $\mathbb{P}_b = \mathbb{P} - \mathbb{P}_a + p\mathbb{I}$ which vanishes as long as $\epsilon = 0$. Computing the steady state stress $\mathbb{P}_b^\infty$ after the osmotic shock we can compute the associated pressure $p_b^\infty = -\mathrm{tr}(\mathbb{P}_b^\infty)/3 = -K_d\mathrm{tr}\epsilon^\infty$. This pressure field is experimentally reported in Fig 4b of [9] and depends on the three non-dimensional parameters $R_c/R_m$, $p_a/G$ and $\beta$. For simplicity, we fix $\beta = -1$ and fit the two remaining parameters $R_c/R_m \simeq 0.1 \in (0, 0.3)$ and $p_a/G \simeq -0.7 \in (-1.1, -0.3)$ with the data of [9]. The parenthesis denote the 95% confidence interval of the fit. See Fig. 7 (a) for the experimental data overlaid with the fit. With our AFM measurements of MCS compression, we can estimate the shear modulus of the MCS $G \simeq 280 \in (270, 290)$ Pa (See Section. 3) leading to a realistic magnitude of the active stress $|p_a| \simeq 200$ Pa.

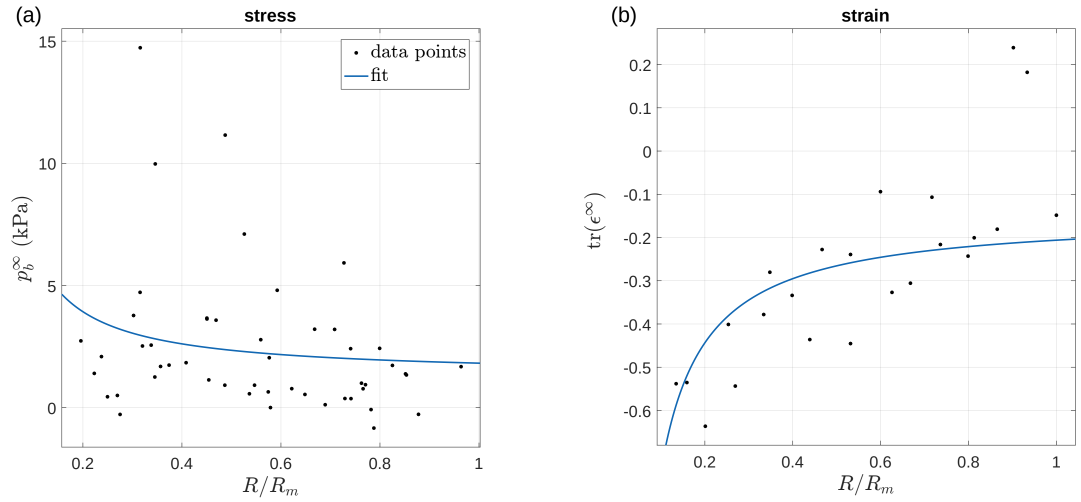


Figure 7: (a) Steady state pressure $p_b^\infty$ within the MCS under a $\Pi_e = 5$kPa osmotic compression (data from [9]). The full line corresponds to the analytic expression $p_b^\infty$ with parameters $\beta = -1$, $R_c/R_m = 0.1$ and $p_a/G = -0.7$. (b) Steady state volumetric strain estimated from the variation of the nucleus distance (data from [7]). The parameters are the same as (a) and the only fitted parameter is the drained modulus $K_d = 8.8$ kPa.

Interestingly, our expression of $p_b^\infty$ gives a plausible explanation for the "pressure jump" reported in [9] between the value of $p_b^\infty(R_m) = \Pi_e + \Pi_e p_a/G(1 - (R_m/R_c)^\beta)$ at the MCS surface and the osmotic pressure $\Pi_e$. Note that $\Pi_e - p_b^\infty(R_m) \geq 0$ vanishes in the absence of active stress ($p_a = 0$) showing that this jump is of active origin. In the limit where $R_c \ll R_m$, the pressure jump takes a simple form $\Pi_e - p_b^\infty(R_m) = -\Pi_e p_a/G$ which can be used to estimate the active pressure.

Finally, we can use independent measurements of the volumetric strain within the MCS (see [7]) to estimate the bulk modulus $K_d$. Using the best fit parameters from the stress experiment, we fit the only remaining parameter $K_d$ in our analytic expression (13) to obtain $K_d \simeq 8.8 \in (7, 11)$ kPa. This fit is overlaid with the data on Fig. 7 (b).

**Computation of the effective compression modulus.** Based on the displacement field (13), we compute the total relative loss of volume,

$$\frac{\Delta V}{V} = \frac{1}{|\Omega|}\int_\Omega \mathrm{tr}\epsilon^\infty d\underline{X} = \frac{3}{R_m^3}\int_0^\infty R^2 \mathrm{tr}\epsilon^\infty dR = -\frac{\Pi_e}{K_d}\left(1 - \frac{p_a\left(3\left(\frac{R_m}{R_c}\right)^\beta - \beta - 3\right)}{(\beta+3)G}\right). \quad (14)$$

The second term on the right hand side of (14) results in an active compression modulus which can be attributed to MCS volumetric growth. Indeed, when the MCS reaches its steady state radius $R_c = (3/(3+\beta))^{1/\beta} R_m$, the active contribution vanishes and the compression modulus reduces to $K_d$.

Expression (14) can be further simplified to,

$$\frac{\Delta V}{V} = -\frac{\Pi_e}{K_d}\left(1 + \frac{p_a}{G}\right)$$

when $R_c \ll R_m$ and results in the active compression modulus

$$K_{\text{active}} = \frac{K_d}{1 + p_a/G}.$$

which is higher than in the passive case. With the above estimates of $K_d$ and $p_a/G$, we obtain $K_{\mathrm{active}} \simeq 29$ kPa which is very close to the value $K_{\mathrm{exp}} \simeq 28 \in (21, 34)$ kPa measured experimentally (See Fig. 8 (a)).

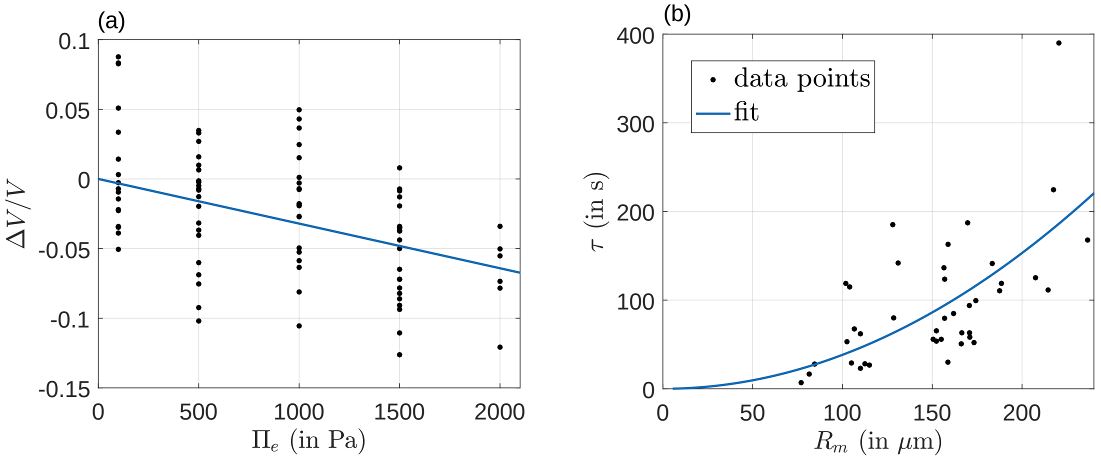


Figure 8: (a) Total volume loss measured when MCS are submitted to a small external osmotic pressure. Dots are experimental data points and the full line is a linear fit with slope $K_{\mathrm{exp}} \simeq 28 \in (21, 34)$ kPa. Experimentally, the volume is deduced by measuring the projected area of MCS, imaged by phase contrast respectively 5 minutes before and 20 minutes after the osmotic shock. This timescale is short as compared to the cell division time (18 h for CT26 cells). (b) Relaxation time of the MCS to its steady radius after an osmotic shock as a function of the MCS initial radius. The fit corresponds to formula (18) with a diffusion coefficient $D_{\mathrm{passive}} \simeq 1.4 \in (1.2, 1.7) \times 10^{-12} \mu\mathrm{m}^2\mathrm{s}^{-1}$ and $p_a/G = -0.7$.

**Water relaxation.** The active poroelastic theory (10) also enables us to predict the dynamics of the water percolation in the MCS pores after the application of an osmotic pressure. Indeed, we have,

$$\frac{\partial \delta n}{\partial t} - \frac{\kappa}{\mu} \nabla^2 p = 0 \tag{15}$$

where the porosity variation following the increase of pore pressure is $\delta n = \mathrm{tr}(\mathbb{F}_a^T \epsilon \mathbb{F}_a)$ and

$$\mathrm{div}(\epsilon \mathbb{P}_a + \det \mathbb{F}_a^{-1} \mathbb{F}_a \mathbb{S}(\mathbb{F}_a^T \epsilon \mathbb{F}_a) \mathbb{F}_a^T) = \nabla p.$$

As in the steady state, this last equation relating the strain and the pore pressure can be explicitly solved in power series of $\delta_a$, $u(R,t) = u_0(R,t) + \delta_a u_1(R,t)$ in the limit $p_a \ll G \ll K_d$ to obtain the unsteady displacement field as a function of the pressure distribution

$$u_0(R,t) = \frac{1}{R^2 K_d} \int_0^R x^2 p(x,t)\, dx$$

and,

$$u_1(R,t) = \frac{1}{G} \left( C[p(x)]R + R \int_0^R f[x; p(x,t)] dx - \frac{1}{R^2} \int_0^R x^3 f[x; p(x,t)] dx, \right)$$

where the constant $C$ and functional $f$ depend only on parameters $\beta$ and $R_c/R_m$ but are too lengthy to be made explicit here. From these expressions, one can determine the functional expression of pore pressure variation $\delta n$. Because of the presence of some involved integration constants, it is easier to express the derivative of such quantity:

$$\frac{\partial \delta n}{\partial R} = \frac{1}{K_d}\left[\frac{\partial p}{\partial R} - \frac{p_a}{G}\left(\beta\left(\frac{R}{R_c}\right)^{\beta}\frac{p}{R} + \frac{\beta\left(\left(\frac{R}{R_c}\right)^{\beta} - \left(\frac{R_m}{R}\right)^3\right) - 3\left(\frac{R_m}{R}\right)^3\left(1 - \left(\frac{R_m}{R_c}\right)^{\beta}\right)}{\beta+3}\frac{\partial p}{\partial R}\right)\right].$$

It is then clear that if $p_a = 0$, plugging this expression into (15), we recover the passive case when water percolation outside the MCS follows a diffusion process with diffusion coefficient

$$D_{\text{passive}} = \kappa K_d/\mu.$$

In contrast, when $p_a \neq 0$, the percolation of water does not follow a diffusion process in general. However, it does in the special limit when $R_c \ll R_m$ that we will further investigate now. Indeed, in that case, as long as $\beta < 0$, the above expression can be simplified to

$$\frac{\partial \delta n}{\partial R} = \frac{1 + p_a/G(R_m/R)^3}{K_d}\frac{\partial p}{\partial R}$$

at the leading order in $R_c/R_m$. Plugging this expression into (15) and introducing the non-dimensional coordinate $x = R/R_m$ and time $s = t/(R_m^2/D_{\text{passive}})$, we obtain the space dependent diffusion equation,

$$\frac{\partial \delta n(x,s)}{\partial s} = \frac{x^2\left(\left(5p_a/G + 2x^3\right)\partial_x \delta n(x,s) + x\left(p_a/G + x^3\right)\partial_{xx}\delta n(x,s)\right)}{\left(p_a/G + x^3\right)^2}$$

which again needs to be solved by expansion in power series of $p_a/G$: $\delta n(x,s) = \delta n^0(x,s) + \frac{p_a}{G}\delta n^1(x,s)$ leading to,

$$\frac{\partial \delta n^0}{\partial s} - \nabla_x^2 \delta n^0 = 0 \text{ where } \nabla_x^2 . = x^{-2}\frac{\partial}{\partial x}\left(x^2\frac{\partial .}{\partial x}\right) \tag{16}$$

and,

$$\frac{\partial \delta n^1}{\partial s} - \nabla_x^2 \delta n^1 = L(x;\delta n^0) \stackrel{\text{def}}{=} x^{-4}\frac{\partial \delta n^0}{\partial x} - x^{-3}\frac{\partial \delta n^0}{\partial xx}. \tag{17}$$

To estimate the time needed for the water to percolate outside the MCS, we consider the above system with $n^0(x,0)$ a Dirac mass at the origin such that all the water is concentrated there. Then solving (16) we obtain a normal distribution for $\delta n^0$,

$$\delta n^0(x,s) = \frac{e^{-\frac{x^2}{4s}}}{2\sqrt{\pi}s^{3/2}}$$

Plugging this expression into (17), we can solve for $\delta n^1$. In particular, we can compute the integrals,

$$\int_0^{\infty} x^2 \delta n^0 dx = 1 \text{ and } \int_0^{\infty} x^2 \delta n^1 dx = \frac{1}{6\sqrt{\pi}s^{3/2}}.$$

The average square distance over which water propagates is

$$\langle X^2 \rangle \stackrel{\text{def}}{=} \frac{\int_0^{\infty} x^4 \delta n dx}{\int_0^{\infty} x^2 \delta n dx}.$$

As, using (16) and (17) and Green's formula we have

$$\begin{aligned}\int_0^\infty x^4 \partial_s \delta n dx &= \int_0^\infty x^4 \left( \nabla_x^2 \delta n^0 + \frac{p_a}{G} \nabla_x^2 \delta n^1 + \frac{p_a}{G} L(x;\delta n^0) \right) dx \\ &= 6 \int_0^\infty x^2 \delta n^0 dx + \frac{6 p_a}{G} \int_0^\infty x^2 \delta n^1 dx + \frac{p_a}{G} \int_0^\infty x^4 L(x;\delta n^0) dx \\ &= 6,\end{aligned}$$

we obtain,

$$\frac{d\langle X^2\rangle}{ds} = \frac{\int_0^\infty x^4 \partial_s \delta n dx}{\int_0^\infty x^2 \delta n dx} - \frac{\int_0^\infty x^2 \partial_s \delta n dx}{\left(\int_0^\infty x^2 \delta n dx\right)^2} \int_0^\infty x^4 \delta n dx.$$

Thus, at first order in $p_a/G$ we obtain,

$$\frac{d\langle X^2\rangle}{ds} = 6 + \frac{p_a}{2G\sqrt{\pi} s^{3/2}} \text{ and } \langle X^2\rangle = 6s - \frac{p_a}{G\sqrt{\pi s}}.$$

The above computation therefore shows that the active stress contributes to the dynamics of the water percolation out of the MCS as we finally obtain a first order correction of the classical Stokes-Einstein result $\langle X^2\rangle = 6s$. In dimensional form this implies that the effective time of water percolation $\tau$ satisfies,

$$R_m^2 = 6\tau D_{\text{passive}} - \frac{p_a R_m^3}{G\sqrt{\pi\tau D_{\text{passive}}}},$$

such that at first order,

$$\tau = \frac{R_m^2}{6D_{\text{passive}}} \left( 1 + \frac{p_a}{G}\sqrt{\frac{6}{\pi}} \right). \tag{18}$$

As $p_a$ is negative, this implies that activity speeds up the water percolation in the MCS during compression leading to an active diffusion coefficient,

$$D_{\text{active}} = \frac{D_{\text{passive}}}{1 + \frac{p_a}{G}\sqrt{\frac{6}{\pi}}}.$$

However, this linear approximation clearly fails for values of $p_a/G < -\sqrt{\pi/6} \simeq -0.72$ as it leads to a negative percolation duration. Thus we only use this formula to obtain an upper bound on $D_{\text{passive}}$. Since the relaxation time $\tau$ is experimentally measured as a function of the MCS radius $R_m$, using (18), we can fit the value of $D_{\text{active}} \simeq 4.4 \in (4, 4.8) \times 10^{-11} \mu\text{m}^2\text{s}^{-1}$. See Fig. 8 (b). Taking into account the error associated to $p_a/G = -0.7 \in (-1.1, -0.3)$, we obtain the upper bound for the $D_{\text{passive}} < 2.8 \times 10^{-11} \mu\text{m}^2\text{s}^{-1}$.

With this measurement of $D_{\text{passive}}$, we can estimate an upper bound of the MCS permeability $\kappa = \mu D_{\text{passive}}/K_b \lesssim 3.2 \times 10^{-18}$ m$^2$ which is comparable to previous measurements [20].

#### 4.4 ECM rheology and volume exclusion of cells can explain the MCS mechanical properties

In this section, we demonstrate that the effective rheological coefficients of the MCS, $K_d$ and $\kappa$ can be interpreted as stemming from the ECM while cells are considered as impermeable incompressible objects which are only responsible for volume exclusion.

**MG as an ECM proxy** As interstitial ECM is difficult to access, we use MG balls to estimate the rheological properties of ECM. Balls of MG are typically passive poro-elastic materials to which we can apply results of Section 4.2.

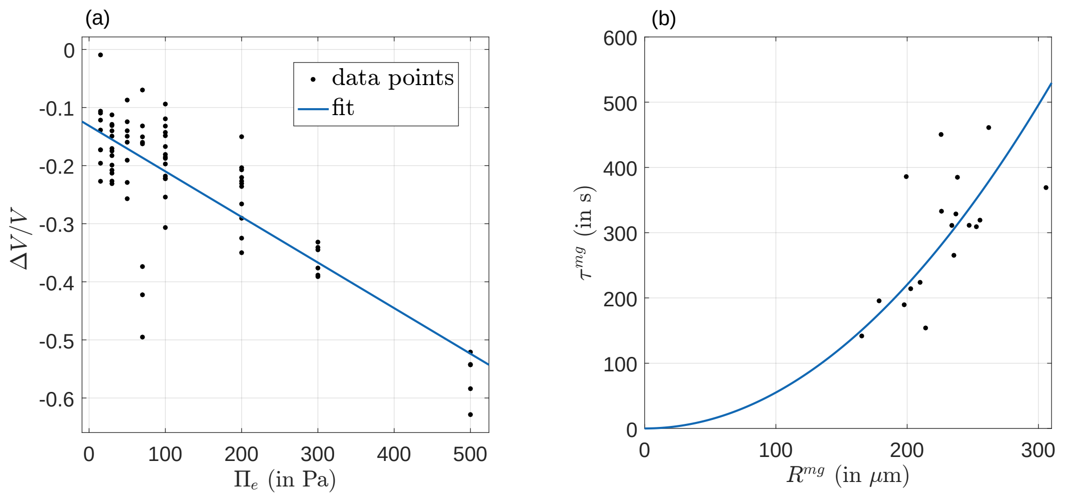


Figure 9: (a) Compression of MG balls as a function of osmotic pressure. The linear fit is $\Delta V/V = -\Pi_e/K_d^{\mathrm{mg}} + \mathrm{offset}$ where $K_d^{\mathrm{mg}} \simeq 1270 \in (1060, 1490)$ Pa and offset $\simeq -0.13 \in (-0.1, -0.15)$. (b) Relaxation time of MG balls as a function of their initial radius. The quadratic fit corresponds to $\tau^{\mathrm{mg}} = (R^{\mathrm{mg}})^2/(6D^{\mathrm{mg}})$ where $D^{\mathrm{mg}} \simeq 3 \in (2.6, 3.4) \times 10^{-11}\mathrm{m}^2\mathrm{s}^{-1}$.

**Drained modulus** We thus begin by estimating the bulk modulus $K_d^{\mathrm{mg}}$ of MG balls by applying osmotic pressure shocks to them. See Fig. 9 (a). By fitting the data with a linear curve, we obtain $K_d^{\mathrm{mg}} \simeq 1270 \in (1060, 1490)$ Pa. Note the presence of an offset in the compression data probably due to some ionic affinities of some MG components that occur when the Dextran solution used to perform the osmotic shock is introduced. To complement this measurement, we also perform AFM compression of MG layers to estimate their shear modulus $G^{\mathrm{mg}} \simeq 100$ Pa (See Section. 3) which is indeed negligible compared to the bulk modulus.

If we now suppose that cells within the MCS are incompressible inclusions (their bulk modulus is $\sim 800$ kPa $\gg K_d^{\mathrm{mg}}$ [17]) coated by MG, we can deduce the bulk modulus of the MCS from the Hashin-Shtrikman upper bound [14, 13] which we expect to be a good approximation [16]

$$K_d \simeq K_d^{\mathrm{mg}} + \frac{1-n_m}{n_m}(K_d^{\mathrm{mg}} + 4G^{\mathrm{mg}}/3) \simeq \frac{K_d^{\mathrm{mg}}}{n_m},$$

where $n_m$ is the average MCS porosity prior to osmotic compression. The porosity $n_m \simeq 0.14$ was estimated in Section. 1.1. We obtain an estimate of $K_d \simeq 9$ kPa which is very close to the one suggested by our theory and experimental measurements presented above.

**Permeability** As in the MCS case, we can also measure the characteristic time associated to water percolation needed to compress MG balls to estimate the diffusion coefficient $D^{\mathrm{mg}} \simeq 3 \in (2.6, 3.4) \times 10^{-11}\mathrm{m}^2\mathrm{s}^{-1}$. See Fig. 9 (b).

Since $\kappa^{\mathrm{mg}} = \mu D^{\mathrm{mg}}/K_d^{\mathrm{mg}}$, we estimate the MG permeability to be $\kappa^{\mathrm{mg}} = 2.3 \times 10^{-17}\mathrm{m}^2$. This value is of the correct of order of magnitude as permeability scales with the square of the characteristic mesh size which is estimated to be roughly 10 nm (See Fig. 3).

Again, considering the cells as almost impermeable compared to the extra-cellular space, we can estimate the MCS permeability according to the classical Maxwell result as,

$$\kappa = \frac{2n_m}{3-n_m}\kappa^{\mathrm{mg}} \simeq 2.3 \times 10^{-18}\mathrm{m}^2$$

which is comparable to the previously computed upper bound $3.2 \times 10^{-18}$ $\mathrm{m}^2$.

##### 4.5 Conclusion

We have first presented an effective (single medium) active poro-elastic theory to model the short timescale response of a MCS under osmotic compression. Comparing such theory with experimental results, we have estimated two key passive phenomenological coefficients that control the compression: the drained bulk modulus of the MCS and its permeability.

From a mechanical perspective the MCS can be considered as a composite material made of cells and ECM and we have shown that both the effective drained modulus and the permeability can be attributed to ECM properties corrected by volume exclusion of incompressible and impermeable cells.

[6] Olivier Coussy. *Poromechanics*. John Wiley & Sons, 2004.

[7] Morgan Delarue, Jean-François Joanny, Frank Jülicher, and Jacques Prost. Stress distributions and cell flows in a growing cell aggregate. *Interface focus*, 4(6):20140033, 2014.

[8] Morgan Delarue, Fabien Montel, Ouriel Caen, Jens Elgeti, Jean-Michel Siaugue, Danijela Vignjevic, Jacques Prost, Jean-François Joanny, and Giovanni Cappello. Mechanical control of cell flow in multicellular spheroids. *Physical review letters*, 110(13):138103, 2013.

[9] ME Dolega, Morgan Delarue, François Ingremeau, Jacques Prost, Antoine Delon, and Giovanni Cappello. Cell-like pressure sensors reveal increase of mechanical stress towards the core of multicellular spheroids under compression. *Nature communications*, 8:14056, 2017.

[10] Massimiliano Fraldi and Angelo R Carotenuto. Cells competition in tumor growth poroelasticity. *Journal of the Mechanics and Physics of Solids*, 112:345–367, 2018.

[11] Alain Goriely. *The mathematics and mechanics of biological growth*, volume 45. Springer, 2017.

[12] Karine Guevorkian, Marie Josée Colbert, Mélanie Durth, Sylvie Dufour, and Françoise Brochard-Wyart. Aspiration of biological viscoelastic drops. *Physical Review Letters*, 104(21):1–4, 2010.

[13] Zvi Hashin. The elastic moduli of heterogeneous materials. *Journal of Applied Mechanics*, 29(1):143–150, 1962.

[14] Zvi Hashin and Shmuel Shtrikman. A variational approach to the theory of the elastic behaviour of multiphase materials. *Journal of the Mechanics and Physics of Solids*, 11(2):127–140, 1963.

[15] Ora Kedem and Aharon Katchalsky. Thermodynamic analysis of the permeability of biological membranes to non-electrolytes. *Biochimica et biophysica Acta*, 27:229–246, 1958.

[16] Graeme W Milton. The theory of composites. *Cambridge University Press*, 2002.
